# Timing and magnitude of climate driven range shifts in transboundary fish stocks challenge their management

**DOI:** 10.1101/2021.08.26.456854

**Authors:** Juliano Palacios-Abrantes, Thomas L. Frölicher, Gabriel Reygondeau, U. Rashid Sumaila, Alessandro Tagliabue, Colette C.C. Wabnitz, William W.L. Cheung

## Abstract

Climate change is shifting the distribution of shared fish stocks between neighboring countries’ Exclusive Economic Zones (EEZ) and the high seas. The timescale of these transboundary shifts determines how climate change will affect international fisheries governance. Coupling a large ensemble simulation of an Earth system model to a species distribution model, we show that by 2030, 23% of transboundary stocks would have shifted and 78% of the world EEZs will experience at least one shifting stock under a high emission climate change scenario. By the end of this century, 81% of EEZ waters will see at least one shifting stock with a total of 45% of stocks shifting globally, under a high emissions scenario. Importantly, many countries that are highly dependent on fisheries for income, food and nutrition security, as well as livelihoods emerge as hotspots for transboundary shifts showing early, and sometimes past shifts. Existing fisheries agreements need to be assessed for their capacity in addressing transboundary shifts, and strengthened where necessary to limit conflict over these fish stocks while new agreements are urged to considere this problematic in order to be resilient to global change.

## Introduction

Over the past century, human activities have dramatically altered the physical and biogeochemical conditions of the ocean, resulting in warmer, more acidic and less oxygenated waters (IPCC, 2019). Marine species’ distributions reflect species’ preferences for discrete environmental conditions (Hutchinson, 1957). As a result of changing ocean conditions, many marine species are shifting their distributions towards higher latitudes, deeper water or following local temperature gradients to remain within their optimal environmental “niche” (Poloczanska et al., 2016). The biogeography of marine species is projected to continue to shift as ocean conditions change through the 21st century (Cheung et al., 2010), impacting fish stocks, fisheries production and economies (Sumaila et al., 2019). These impacts are compromising our capacity to reach international sustainability goals under the United Nations (UN) 2030 Agenda for Sustainable Development such as Goal 14 - life below water (Pecl et al., 2017; Singh et al., 2017; United Nations, 2018). The projected risks and impacts can be reduced by improving the effectiveness of current governance and fisheries management, including for species that cross international borders, also known as ‘shared stocks’ (Pinsky et al., 2018).

The concept of shared stocks was developed following the ratification of the UN on the Law of the Sea (UNCLOS) and the claiming of Exclusive Economic Zones (EEZs) by Coastal States (United Nations, 1986). As defined by the UN’s Food and Agriculture Organization, shared stocks can be classified into four non-exclusive categories: (*i*) transboundary stocks, which cross neighboring EEZs; (*ii*) straddling stocks that, in addition to neighboring EEZs, also visit the adjacent high seas; (*iii*) highly migratory stocks, mainly tunas and bill-fishes, that migrate across vast oceanic regions including both the high seas and EEZs; and (*iv*) discrete stocks that are only present on the high seas (G. Munro et al., 2004). This study focuses on transboundary stocks exploited by fisheries operating within EEZs. Under UNCLOS, countries are responsible for the management of stocks within their EEZs and encouraged to cooperate when stocks are shared (United Nations, 1986). A recent study estimates that there are 633 transboundary fish species globally, representing 67% of identified exploited taxa. Between 2005 and 2010, these species yielded an annual average of 48 million tonnes of catch and USD 78 billion in fishing revenue (Palacios-Abrantes, Reygondeau, et al., 2020).

Most fisheries management is poorly adapted to address species’ shifting distributions because of changing ocean conditions. The effectiveness of fisheries management for transboundary species is challenged by shifts in stocks’ distribution under climate change (Oremus et al., 2020; Pinsky et al., 2018; Pinsky et al., 2020; Pinsky & Mantua, 2014). In most cases, catch or fishing effort quotas for transboundary stocks are based on historical records and do not necessarily consider the full distribution range of the stock (Fredston-Hermann et al., 2018), nor the effects of a changing climate on fish stocks and associated fisheries (Palacios-Abrantes, Reygondeau, et al., 2020; Sumby et al., 2021). While many transboundary stocks experience natural seasonal variation in their distribution (e.g., migration, reproductive cycle), misalignment between fisheries resources’ allocation and distributional shifts beyond such natural variation have previously resulted in unsustainable fishing levels and international disputes (Crespo et al., 2020; Miller et al., 2013; Spijkers & Boonstra, 2017). For example, Pacific salmon species (*Oncorhynchus sp*.), jointly managed by Canada and the U.S., were heavily fished in the early 1990s after a stock shift favored the Alaskan fleet over the Canadian fleet (Miller et al., 2013). Continuing climate change is expected to exacerbate such patterns (Pinsky et al., 2018; Sumaila et al., 2011). Understanding when climate change will affect the sharing dynamics of transboundary stocks and the intensity of the resulting impacts is important for developing climate resilient international ocean governance and achieving the goals set out under the 2030 Agenda (Link et al., 2010; Pinsky et al., 2018; Sumaila et al., 2020).

Here, we employ a mechanistic population dynamic model driven by output from a comprehensive Earth system model with ten ensemble members to project transboundary stocks’ distribution across the 280 EEZs of 198 coastal countries/political entities under a high greenhouse gas emissions scenario (Methods). We then developed a Transboundary Index to evaluate range shifts in the shared distribution of transboundary stocks. The index is based on changes in the distance of the stock’s abundance centroid and that of the EEZs that share a given stock (Fig. 1). By employing the concept of *time of emergence*, we estimate the year in which the 633 transboundary species (9,132 stocks) are projected to shift their shared distribution beyond historical variations (Fig. 1). Such estimation is based on two thresholds representing a 67 and a 95% probability of shift. Finally, we apply the Game Theory concept of *threat point* (G. R. Munro, 1979 for its application to fisheries; See Sumaila et al., 2020) to quantify the intensity of changes in the shared distribution of transboundary stocks between neighboring EEZs (Methods). This assumes that countries are adapted to the historical sharing, but no country would be willing to collaborate if the future catch proportion of the transboundary stock is lower than it has ever been (Palacios-Abrantes, Sumaila, et al., 2020; Sumaila et al., 2020). We treat each species in an EEZ as a single stock due to the lack of more spatially resolved data to delineate the boundary of a managed population (Palacios-Abrantes, Reygondeau, et al., 2020) and only consider shared stocks between neighboring EEZs (i.e., excluding the high seas).

**Figure 1:**
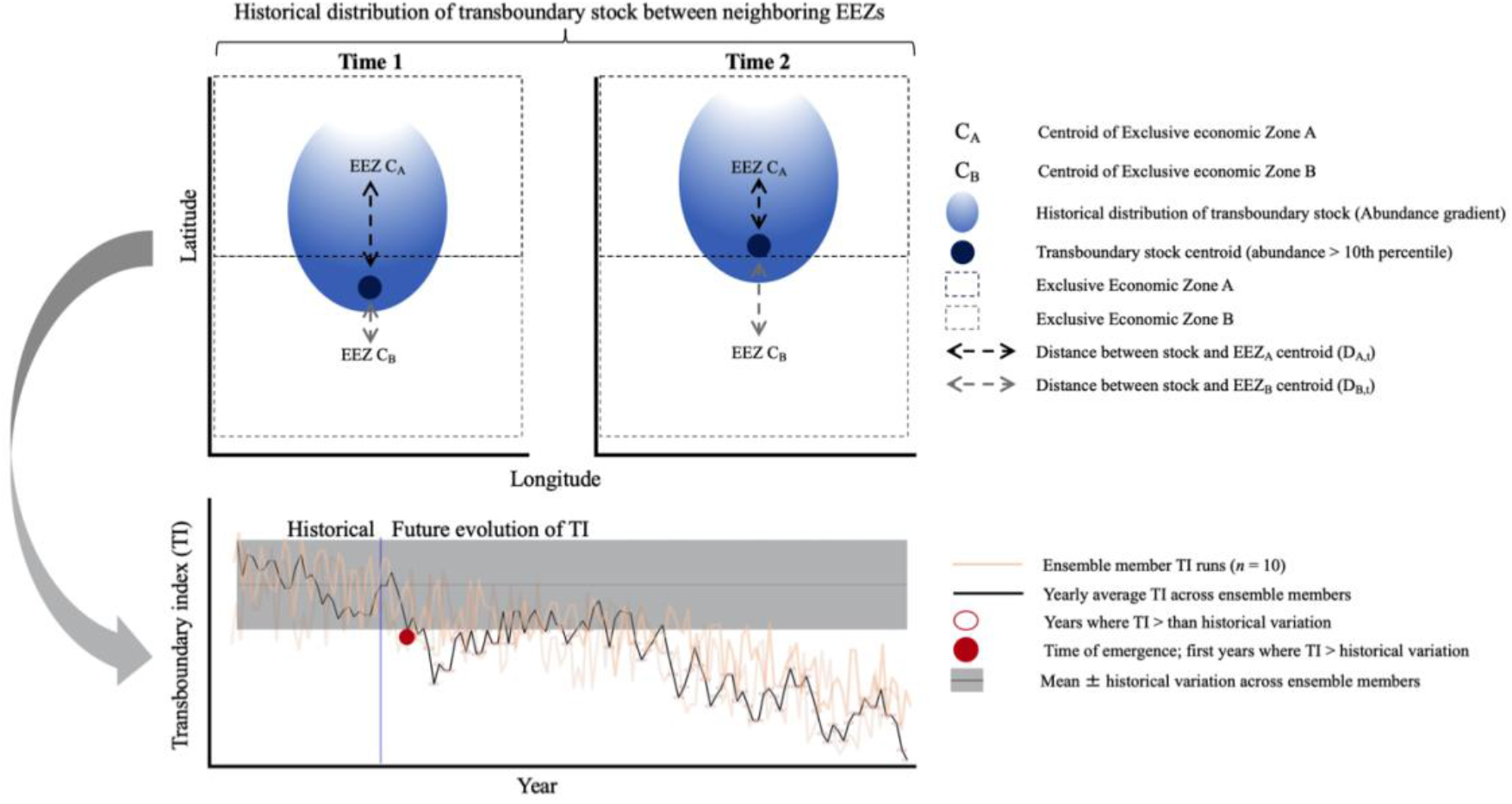
Schematic diagram of the transboundary index (TI) to determine the time of emergence of transboundary stocks from one Exclusive Economic Zone (EEZ) to another. The index is based on the distance between the distributional centroid of the transboundary stock and the geographic centroid of the neighboring EEZ sharing the stock (Top panel). Time of emergence is defined as the first year when the TI overshoots historical values (Bottom panel). Only showing a subset of ensemble members for clarity.

## Materials and Methods

### Databases and species selection

The analyses detailed herein are based on 633 exploited marine transboundary species that account for 80% of the catch taken from the world’s EEZs between 2005 and 2014 (Palacios-Abrantes, Reygondeau, et al., 2020). When a species was shared by a pair of neighboring EEZs it was considered a transboundary stock (Palacios-Abrantes, Reygondeau, et al., 2020; Teh & Sumaila, 2015), resulting in a total of 9,132 transboundary stocks. Using the transboundary species Atlantic cod (*Gadus morhua*) as an example and this study’s definition of transboundary stock, the United States and Canada share a stock of Atlantic cod, and Canada and Greenland share a separate stock of Atlantic cod, but the United States and Greenland do not as they do not have juxtaposing EEZs. We defined the boundaries of the world’s EEZs using the Sea Around Us spatial division (updated 1 July 2015; http://www.seaaroundus.org), noting that it subdivides the EEZs of 198 coastal states into 280 regions (Fig. S1) including island territories. We determined the intersections between polygons using the R package sf (Pebesma et al., 2018). Each EEZ was categorized by geopolitical region according to the United Nations (https://population.un.org/wpp/DefinitionOfRegions/) and biome (Reygondeau, 2019). The habitat preference of each species was determined following FishBase (http://www.fishbase.org) for fish species and SeaLifeBase (http://www.fishbase.org) for invertebrates (Table 1). For each stock and EEZ, we used Sea Around Us data to estimate catch and fishing revenue from fishing activities. We report both average catch and revenue for the last available decade (2005-2014) (Sumaila et al., 2015; Tai et al., 2017; Zeller et al., 2016).

### Projecting species distributions under climate change

We projected the future distribution of transboundary stocks using a dynamic bioclimatic envelope model (hereafter called DBEM). The DBEM determines the environmental space a species will occupy based on physiology, habitat suitability, depth and latitudinal ranges, and spatial population dynamics as well as preferences for sea temperature, salinity, oxygen content, sea ice extent (for polar species) and bathymetry. The DBEM then estimates species’ abundance and maximum catch potential (a proxy for maximum sustainable yield) over a regular spatial grid of 0.5 of latitude per 0.5 of longitude. Below we provide an overall description of the model. Please see Cheung et al. (2010); Cheung et al. (2009); Cheung et al. (2016) for a more detailed version. Importantly, the DBEM is able to project catches by Exclusive Economic Zones that are consistent with observational-based estimates of catch from 1950 to 2016 (Cheung et al., 2016, Cheung et al. *in revision*). For each species, the DBEM uses the Sea Around Us (www.searaundus.org) distributional data and environmental variables from 1970 to 2000 to estimate the current distribution and environmental profile of each species. Moreover, it projects the future habitat suitability in each grid-cell (*i*) based on the following components;

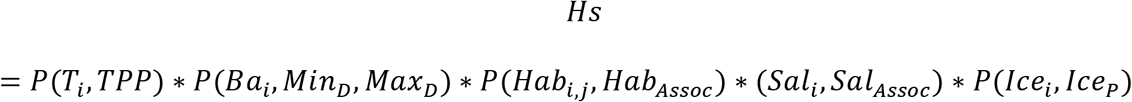

where *T* is seawater temperature, *TPP* is the species temperature profile, *Ba* is bathymetry, *Min_D_* and *MaxD* are the species minimum and maximum depth limits, *Hab* is the proportion of area of the habitat type *j* relative to the total seawater area of the cell *i*, *Hab_Assoc_* is an habitat association index, *Sal* is the salinity class of cell *i* and *Sal_Assoc_* is a salinity class association index. Finally, for polar regions and species, *Ice* is sea ice extent, and *Ice_P_* is association to sea ice. For pelagic species, the model uses environmental variables at the surface whereas demersal and benthic species’ distributions are driven by ocean bottom variables.

The DBEM also integrates each species preference profile with physiology and population dynamics to project relative biomass assuming that spatio-temporal dynamics are determined by intrinsic growth rate, larval dispersion, and adult migration;

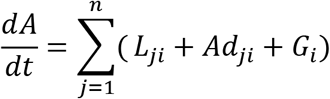

where *A_i_* is the relative abundance of the gri-cell *i*, *G* is the intrinsic population growth and *L_ji_* and *Ad_ji_* are settled larvae and net migrated adults from surrounding cells (*j*), respectively. Larval dispersal is modeled through ocean current with an advection-diffusion-reaction model. Finally, *G_i_* estimates the intrinsic growth following a logistic equation and the intrinsic rate of population increase (*r*):

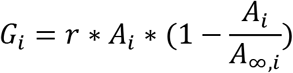

Finally, the DBEM projects maximum catch potential by setting fishing mortality to half of the intrinsic population growth rate of the species (*f* = *r/2*).

The DBEM was driven by simulated ocean conditions from a ten ensemble member simulation of the Geophysical Fluid Dynamics Laboratory Earth system model (GFDL-ESM2M) to project the distributions of the 633 species from 1951 to 2100 (John et al., 2012, 2013; Rodgers et al., 2015). The GFDL-ESM2M was run under historical forcing until 2005 and follows a high greenhouse gas emissions scenario, Representative Concentration Pathway 8.5 (RCP 8.5), over the 2006-2100 period (Riahi et al., 2011). We chose the 1951-2100 time period to match the historical period in the GFDL ESM2M simulations over which the model was forced with observation-based greenhouse gas, aerosol and natural external forcing (Rodgers et al., 2015). Because the main approach of this paper relies on understanding the spatial and temporal variation of a stock’s distribution, we have to understand distribution variability during both historical and future periods, to infer differences between time frames. To this end, each of the ten GFDL-ESM2M ensembles were started from infinitesimally small differences in Earth system initial conditions in 1950, resulting in a unique atmosphere and ocean state at each point in time after about three years for surface and eight years for subsurface waters (Frölicher et al., 2020). By design, variations among ensemble members are then solely due to natural internal variability (e.g., different phases of the El Niño Southern Oscillation-ENSO). Thus, for our experiment, each of the ten GFDL-ESM2M ensemble simulations were started from infinitesimally small differences in Earth system initial conditions in the year 1950, resulting in a unique atmosphere and ocean state at each point in time after about three years for surface waters and eight years for subsurface waters (Frölicher et al., 2020).

### Calculating an index of transboundary range shift

We developed a Transboundary Index (TI) to evaluate range shifts in the shared distribution of transboundary stocks under climate change. This index represents the shift in the distribution centroid of a transboundary stock relative to the centroid of neighboring EEZs that share this stock (Fig. 2). The centroid of a transboundary stock was determined by the average (μ) latitude (*lat_ts_*) and longitude (*lon_ts_*) across grid cells within the neighboring EEZs sharing the stock. Therefore,

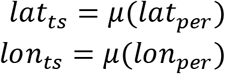

where *lat_per_* and *lon_per_* are the latitudes and longitudes of the grid cells holding the specified percentile (*per^th^*) of the projected transboundary stock abundance. To focus on areas where transboundary stocks are more abundant and fishing activities are more likely to take place, we included grid cells where the projected stock abundance within the neighboring EEZs sharing the stock was above the 95^*th*^ percentile. A sensitivity analysis using a subset of species (n = 34) for all EEZs to examine the effects of different thresholds (per = 20^*th*^, 50^*th*^ and 90^*th*^ percentile) on the calculated index value showed no apparent difference between percentiles (Fig. S2). The centroid of each EEZ was estimated using the st package in R (Fig. S1). For each ensemble member, neighboring EEZs and transboundary stock, we computed the distance between centroids (i.e., EEZ and stock) using the geosphere package in R assuming the Earth is a perfect sphere and ignoring geographic barriers;

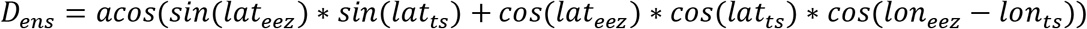

where *lat_eez_* and *lat_ts_* are the latitudes of the EEZ and transboundary stock centroids, respectively, and *lon_eez_* and *lon_ts_* are associated longitudes. Then, for each year and for each ensemble member between 1951 and 2100 we calculated the transboundary index (TI) as follows:

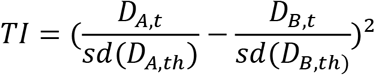

where *D_A_* and *D_B_* represent the distance between the distribution centroids of a stock and the centroid of the neighboring EEZs (A and B) sharing the stock for each time step from 2006 to 2100 (*t*); and *sd* is the standard deviation of the historical (th, 1951 - 2005) centroid distribution for *D_A_* and *D_B_*. The TI was smoothed to a 10-year average to reduce interannual variability (Frölicher et al., 2016) and to match transboundary fisheries management that often integrates a longer time-frame in their consideration.

**Figure 2:**
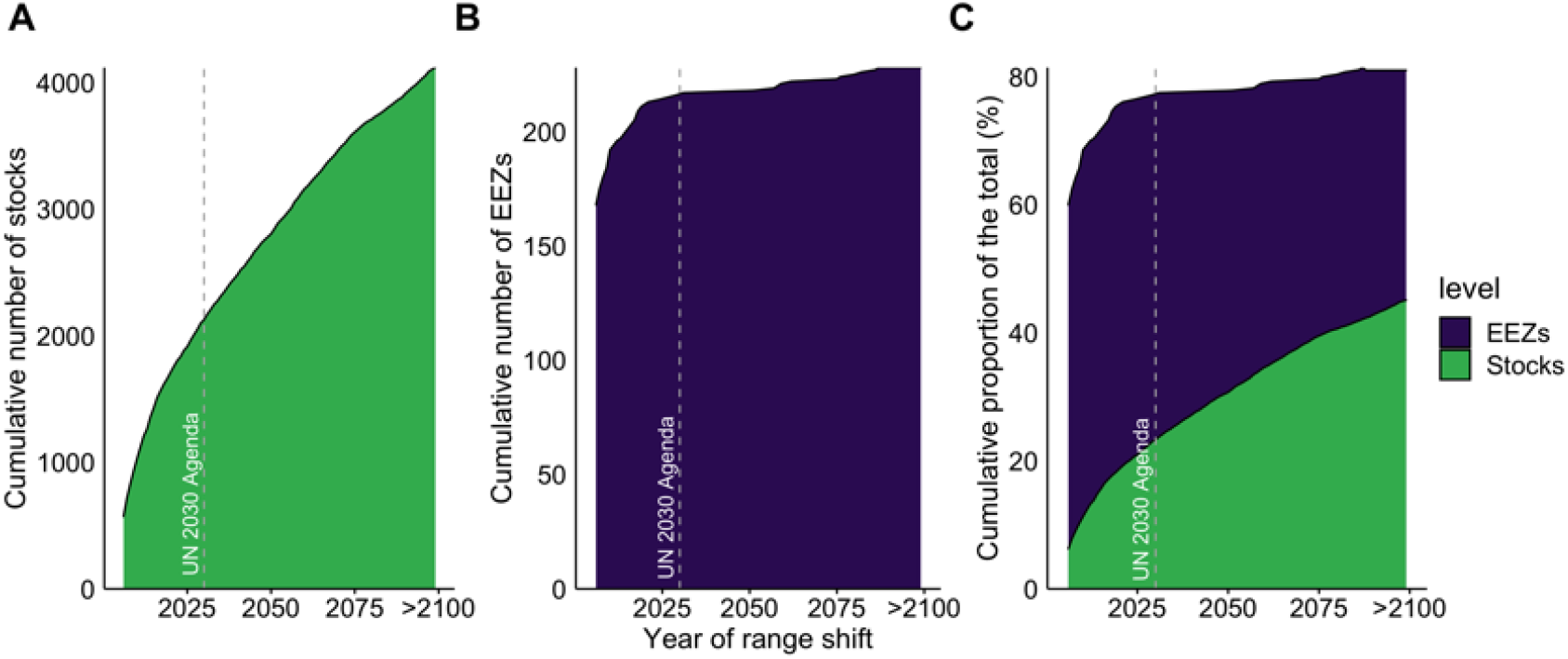
Timeline of shifts in transboundary shared stock distribution. (A) Number of stocks (cumulative) whose distribution centroids that have shifted beyond natural variations over time (from 2006 to >2100), (B) Number of EEZs (cumulative) with the distribution centroid of at least one stock that have shifted beyond natural variations and (C) proportion of total shifting stocks and EEZs with at least one stock shifting beyond natural variability. Dashed lines represent the year by which countries have committed to reach full implementation of the 2030 Agenda, as well as mid and late change in range shift (Methods), respectively.

### Calculating shifts in the shared distribution of transboundary stocks

Knowing the point in time at which the distribution of a shared stock will diverge from historical variability is important to inform decision makers by when, ideally, climate adaptation plans will need to have been implemented (Link et al., 2010). To that end, we adopted the concept of time of emergence, commonly applied to multiple oceanic physical and biogeochemical variables (Cheung & Frölicher, 2020; Henson et al., 2017; Keller et al., 2014; Rodgers et al., 2015; Schlunegger et al., 2019, 2020) and defined as the moment in time when a signal (e.g., future anthropogenic trend) emerges from the background noise of natural variability (i.e., historical natural variation) (Hawkins & Sutton, 2012). The premise behind time of emergence is that we can only be confident that a significant change has been detected when the signal of anthropogenic climate change is larger than the background noise of natural climate variability (Hawkins & Sutton, 2012). The TI’s natural internal variability was estimated from the ensemble members by firstly averaging the TI from 1951 to 2005 and then estimating the variation (*TI_σ_*) across ensemble members. The TI signal represents the average TI across ensemble members from 2006 to 2100 (*TI_μ_*). We then set a threshold (*tresh*) of *TI_σ_* to define the time of emergence of transboundary stocks (See Fig. 2 for a graphical description):

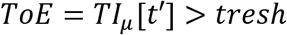

This way we define a shift in a stock’s shared distribution (i.e., time of emergence) as the first year the TI “emerges” from historical internal variability (*tresh*). It assumes that, from a fisheries’ perspective, the first year a stock’s distribution shift from a 10-year average is sufficient to call for caution. Two different arbitrary values of *tresh* were used. A conservative value (*tresh* = 2\**TI_σ_*) representing a 95% probability that the index has emerged from historical variability and a more relaxed value (*tresh* = *TI_σ_*) for a 67% probability.

Since the ToE method is sensitive to the noise estimate (i.e., historical natural variability), we conducted sensitivity analysis where we first calculated the average across the 10 ensemble members for each year between 1951 to 2005 and the calculating the standard deviation over 1950-2005. Under this method, the natural variability is slightly larger resulting in a latter overall time of emergence (average ToE = 2057 vs 2036 of the original method). However, this method projects 1,139 more emerging stocks than the original (5,258 stocks vs 4,119 of the original method). The minimum and maximum ToE remains the same, and the number of EEZs with emerging stocks slightly increases with the alternative method. Further research looking at different approaches of estimating the natural variability component could help reduce the uncertainty in the estimation of the time of emergence of transboundary stocks.

### Quantifying the intensity of transboundary stocks’ range shift

Here, we adopt the concept of threat point to quantify the intensity of changes in the shared distribution of transboundary stocks between neighboring EEZs. The concept of threat point comes from Game Theory and has been widely used in shared fisheries management (Bailey et al., 2010; Clark & Munro, 1975; G. R. Munro, 1979; Sumaila, 2013; Sumaila et al., 2020). In a game theoretic model, a player’s strategy (e.g., to act cooperatively or not) will have direct consequences for other players, which, in turn, will affect the overall outcome of the game (Bailey et al., 2010). Cooperation will often result in the maximization of benefits for the system rather than for individual players. However, for a cooperative strategy to work, the benefit a player gets must not be less than the benefit under a non-cooperative strategy. Thus, the “threat point” is defined as the minimum payoff a player is willing to receive to cooperate in a game-theoretic model (Nash, 1953). We define the threat point as the minimum required catch proportion of a transboundary stock within an EEZ for a country to engage in cooperative management with their sharing neighbor (Palacios-Abrantes, Sumaila, et al., 2020; Sumaila et al., 2020). Any proportion below the defined threat point would result in the unilateral management of the stock assuming that no country will be willing to engage in cooperative management if the future catch proportion of the transboundary stock is lower than it has ever been.

First, we estimated the catch proportion as the stock share ratio (SSR) catch of each transboundary stock shared by neighboring nations between 1951 and 2100. We did this by aggregating the projected catch within the 0.5° x 0.5° grid cells in which the stock is present across neighboring EEZs, and then calculated the proportion of the stock’s catch held within each EEZ (Palacios-Abrantes, Reygondeau, et al., 2020). Second, to reduce the effects of variability, we averaged the calculated proportion of stock occupying a given EEZ into three time periods: the historical time period (*SSR_th_*) from 1951 to 2005, and two future periods, the early 21^*st*^ century, between 2021 and 2040 (*SSR_te_*), and the mid 21^*st*^ century, defined as the average of 2041 to 2060 (*SSR_tm_*). These future time periods were chosen to correspond to the challenges of achieving fisheries-relevant UN-SDGs by 2030 (i.e., the 2030 Agenda), such as SDG 14.4 (end overfishing), SDG 2.4 (ensure sustainable food production systems) and SDG 1.2 (poverty reduction) (Singh et al., 2017; United Nations, 2020). The analysis was replicated for projected stock distributions from each of the ten ensemble members and results were averaged across ensemble members. Third, we defined a threat point for each EEZ’s stock as *SSR_th_* ± 2*σ*, where is the standard deviation of *SSR_th_*. Thus, a change of SSR catch beyond an EEZ’s threat point happens when the future SSR catch exceeds two standard deviations of the historical variations in SSR (i.e., when *SSR_et_* ≧ (*SSR_th_* + 2*σ*) or *SSR_e,m_* ≦ (*SSR_th_* – *σ*). Finally, we estimated the percentage change in SSR (*ΔSSR*) of each future time period (*SSR_tf_*) relative to the historic time period *SSR_th_* for each stock whose share ratio exceeded the threat point following (Palacios-Abrantes, Reygondeau, et al., 2020);

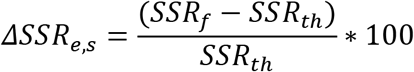

### Statistical analysis

We tested the results for normality (e.g., skewness, kurtosis) and performed two non-parametric Kruskal–Wallis tests by ranks (Hollander & Wolfe, 2013) to investigate geopolitical and ecological differences in the range shift of transboundary species. Specifically, we tested if the habitat preference of transboundary species and the geographic location of EEZs would have any effect on the transboundary stocks distributional range shift. In both cases, our null hypothesis was that there were no significant differences in the year of shift across habitat association or EEZs. All analyses were run using the statistical software R version 3.5.2 (2018-12-20; Eggshell Igloo) with the packages cowplot (Wilke, 2019), data.table (Dowle et al., 2019), geosphere [@], ggrepel (Slowikowski, 2020), gmt (Magnusson, 2017), janitor (Firke et al., 2018), moments (Komsta & Novomestky, 2015), pgirmess (Giraudoux, 2018), rfishbase (Boettiger et al., 2019), sf (Pebesma et al., 2018), sp (Pebesma et al., 2019), tidiverse (Wickham, 2017), tidytext (De Queiroz et al., 2019), viridis (Garnier, 2018), zeallot (Teetor, 2018), and zoo (Zeileis et al., 2019). Code for all analyses is available at https://github.com/jepa/EmergingFish

## Results

### Identifying range shifts in the shared distribution of transboundary stocks

Our results suggest that 4,119 transboundary stocks will experience a range shift beyond historical variability by 2100 (hereafter referred to as ‘shifts’; Fig. 2A), using a two s.d. threshold (i.e., representing a probability of 95% that the stock has shifted). This corresponds to 45% of the studied stocks (Fig. 2C). Using a less conservative threshold (i.e., one s.d. representing a probability of 67% that the stock has shifted) results in 33% (n = 5,745) more shifting stocks, that is, 67% of all studied stocks (Figs. S3 and S4). For both cases, the first shift is modelled to have occurred back in 2006. The average year of shift in the shared distribution of these stocks in all EEZs analyzed is projected to be between 2029 ± 27 years (one s.d. threshold) and 2036 ± 28 years (two s.d. threshold). Furthermore, between 81% and 83% of the world’s EEZs will experience at least one transboundary stock shift by 2100 with 77% to 80% of EEZs recording a distributional stock shift by the UN 2030 Agenda’s deadline (Fig. 2C).

The median year in which transboundary stocks experience a range shift varied significantly according to the species’ habitat association (Methods; Kruskal-Wallis, *X*^2^ = 203.85, DF = 93, *p* < 0.001; using the 2 s.d. threshold; Tables 1, S1 and Fig. S5) and EEZs’ geographic regions (Methods; Kruskal-Wallis, *X*^2^ = 242.11, DF = 93, *p* < 0.001; using the 2 s.d. threshold; Table S3 and Fig. 2A). Overall, most tropical EEZs and stocks will see earlier shifts, with the EEZs of Latin America, the Caribbean and Polynesia experiencing shifts significantly earlier (*p* < 0.05; Tables S1 and S2 than almost any other region (Figs. 3B, S3 and S5). In contrast, EEZs and stocks located in temperate regions, like northern Europe and eastern Asia, are projected to experience much later range shifts. Stocks in a number of EEZs including Brazil and New Zealand’s EEZs are projected to experience no range shifts beyond natural variability by the end of this century (Fig. 3A).

**Figure 3:**
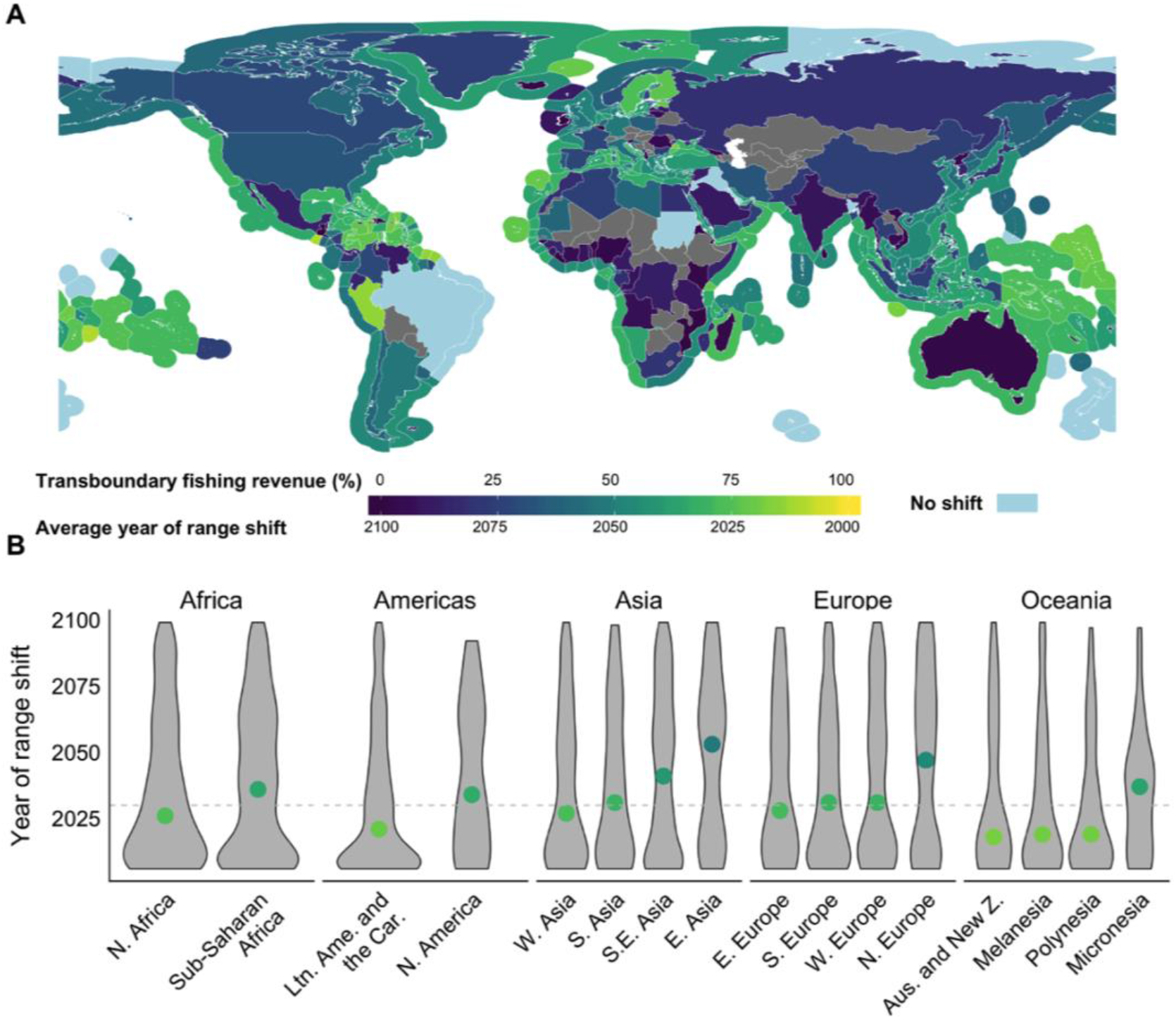
Year of change in the shared distribution of 4,119 transboundary stocks. A) Land polygons show the contribution of shifting stocks to a country or territory’s total fishing revenue from transboundary stocks. Exclusive Economic Zone polygons display average year of range shifts within them. Warm colors are indicative of an early shift/high fishing revenue contribution from transboundary stocks while cool colors represent a late shift/low fishing revenue. EEZs with no distributional shift between 2006 and 2100 are represented in pale blue. B) Year of shared distribution shifts by UN sub-regions. Points color coded by the median year of range shift. Horizontal dashed line represents the year by which countries have committed to reach full implementation of the 2030 Agenda. N = North, S = South, W = West and E = East. Ltn. Ame. and the Car. = Latin America and the Caribbean. Aus and New Z. = Australia and New Zealand.

On average, shifting transboundary stocks represented 27% to 23% (1 and 2 s.d. thresholds) of annual revenue from fisheries targeting transboundary stocks across the world’s EEZs between 2004 and 2010 (Fig. 3A). However, large variation exists, with shifting stocks representing less than 1% of fishing revenue in some countries (e.g., Ireland) and over 90% in others (e.g., Marshall Islands). Moreover, in some EEZs, while few transboundary stocks are projected to shift beyond historical variability, they still represent a large proportion of revenue derived from fisheries within that EEZ (e.g., Peru).

We estimated the current (i.e., 2005 - 2010) contribution of shifting stocks to yearly fishing revenue generated from fisheries targeting transboundary stocks within global EEZs for both thresholds (one and two s.d.). Within five years of the UN 2030 Agenda timeline, 979 transboundary stocks accounting for USD 5.3 billion in fishing revenue are projected to shift within 76 EEZs considering a two s.d. threshold (Fig. 4; above Fisheries >75th percentile). In this case, 321 stocks shifting in just eleven countries account for 4.2 billion USD (Fig. 4). The number of transboundary stocks shifting from historical variability was highest in the EEZs of Spain (n = 122), France (n = 85), China and Portugal (both n = 73), and Senegal (n = 66) (Fig. 4). The values are substantially higher when considering a more relaxed threshold. In this case, 2,136 stocks accounting for USD 24 billion in fishing revenue are projected to shift within five years of the UN 2030 Agenda timeline in 134 EEZs.

**Figure 4:**
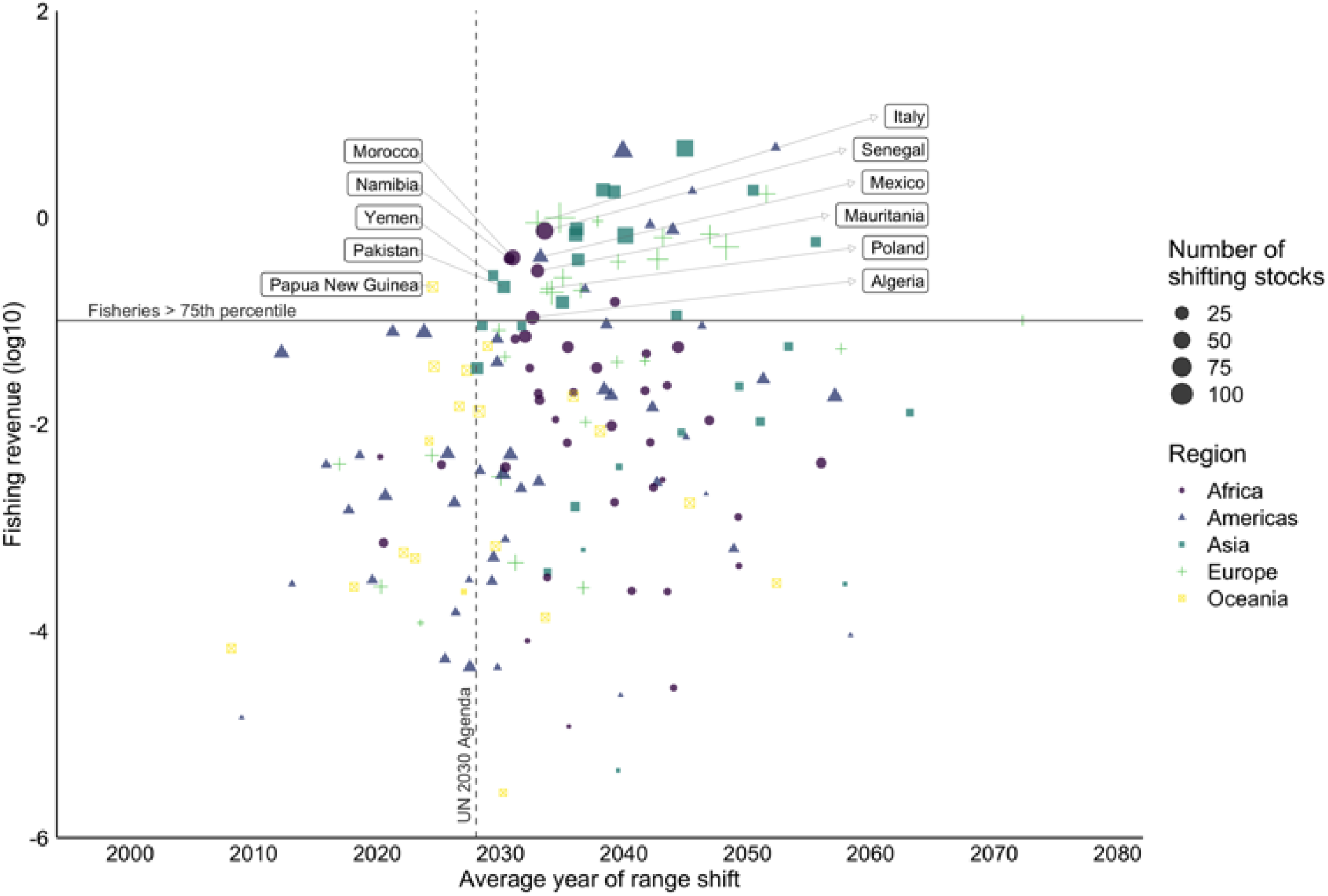
Average range shift of transboundary stocks and corresponding fishing revenue per country. Countries are coded according to different colored symbols representative of 5 distinct regions. The horizontal line separates countries above and below the 75th percentile of annual fishing revenue. Showing country names for the eleven nations above the 75th percentile and projected to have an average shift across transboundary stocks within its EEZ within 5 years of the 2030 Agenda deadline.

### Shifting intensity of transboundary stocks

We estimated the intensity of the distributional shift of transboundary stocks in terms of changes in catch proportion by 2030 (2020-2040) and 2050 (2040-2060), relative to the recent past (1951-2005) (see Methods). Projections show that by 2030, 59% ± 17% of the yearly catch from transboundary stocks will change beyond the historic variability experienced within an EEZ’s (Fig. 5A). Moreover, 85% (n = 239) of the world’s EEZs will have experienced changes in catch proportion of transboundary stocks by 2030. By mid-21st century, the number of EEZs with changing stocks, as well as the number of stocks experiencing changes in catch proportion and the magnitude of that change will increase (Figs. S6 and S7). The direction and intensity of shifts in transboundary stocks are largely related to regional changes in biogeography and the geometry of EEZs (Fig. 5A). Along the Atlantic and Pacific coasts of Northern and Southern America and the Atlantic coast of Southern Africa, shifts in stocks are expected to benefit poleward EEZs. However, shifts along the coasts of Pacific Central America and West Africa occur in an equatorial direction.

**Figure 5:**
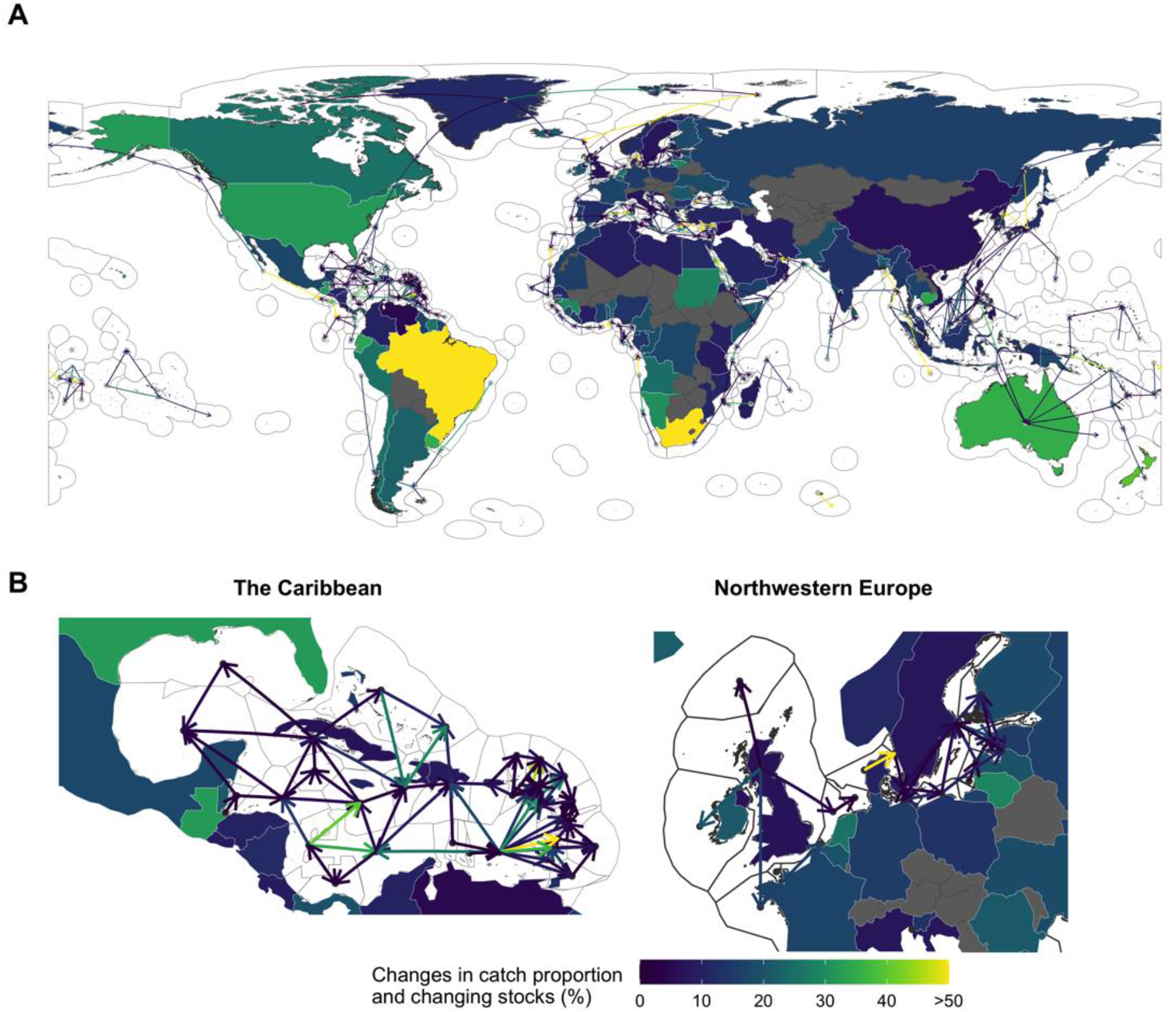
Changes in stock proportion of neighboring Exclusive Economic Zones by 2030 (2021-2040) relative to 1951-2005. Lines represent the average change in transboundary stock share ratio (Methods) with arrows going from EEZ with decreasing stock share (point) to those gaining shares (arrowhead). Land polygons represent the percentage of stocks that are projected to change their stock share ratio (i.e., over two standard deviations from the historical catch proportion) beyond the identified threat point with higher gains identified in warmer colors. Panel B highlights changes for the Caribbean and Northwestern Europe.

## Discussion

### High present-day climate risk for transboundary fisheries

Our findings highlighting early shifts in the shared distribution of transboundary stocks concur with previous studies that have detected changes in marine catch composition and attributed these to climate change (e.g., Cheung et al., 2013; Frainer et al., 2017; Last et al., 2011). For example, in the early 2000s, Humboldt squid (*Dosidicus gigas*) substantially expanded its geographic range poleward, reaching the coast of Washington state (US), in response to climatic, and associated oceanographic and ecological changes (Zeidberg & Robison, 2007). A new fishery targeting Humboldt squid quickly developed on the heels of the species’ range expansion (Pinsky & Mantua, 2014). In the northeast Atlantic, fisheries on Atlantic mackerel (*Scomber scombrus*) are multi-laterally managed by the EU, Norway, Iceland, Russia, and Denmark (on behalf of the Faroe Islands and Greenland) through the North-East Atlantic Fisheries Commission (NEAFC). However, a range expansion of Atlantic mackerel due to environmental variation into Icelandic waters in 2007 resulted in Iceland capturing 6% of the fishery’s total allowable catch and a further 18% in 2008, without consultation with NEAFC, threatening the sustainability of the stock (Spijkers & Boonstra, 2017). These changes resulted in disputes between Iceland and the Faroe Islands, as well as among NEAFC member states (Spijkers & Boonstra, 2017). Other documented cases of early range expansions across international jurisdictions have also been documented for multiple stocks along the European Union regulatory areas (Baudron et al., 2020), the Benguela Current (Potts et al., 2014; Yemane et al., 2014), and the southwest South Atlantic Ocean (Franco et al., 2020). Despite these documented cases, broadly speaking, there is a lack of information and data regarding which and how many transboundary stocks are shifting, where they are shifting to and whether they are jointly managed. This is particularly important for international fisheries management as modeling exercises project that climate change will continue to change the distribution of transboundary stocks to the point that some tropical EEZs will lose them completely (Oremus et al., 2020) while other EEZs, mainly in higher latitudes, will stand to win new stocks (Pinsky et al., 2018).

### Climate risk hotspots for transboundary fisheries management

Our results identify regional “hotspots” of climate risk for transboundary fisheries management that will require the prompt adaptation of management plans (Figs. 3A and 5A). Regions such as the Caribbean are characterized by high levels of warming relative to historical natural variability (Hawkins & Sutton, 2012; IPCC, 2019) and species’ high vulnerability to warming waters (IPCC, 2019). Moreover, they represent relatively low area EEZs that border multiple countries. Game theory predicts that the greater the number of negotiating parties, the harder it is for parties to reach an agreement (Gronbaek et al., 2020). Thus, coordinating the management of shifting transboundary stocks for countries in these regions will be particularly challenging.

Most management plans that are not designed or prepared to respond to range and abundance shifts will be less resilient to climate change (Miller et al., 2013; Sumaila et al., 2020). Focusing on identified “hotspots” will help anticipate any potential increases in and averting fisheries conflicts in coming years. Identified strategies to cope with changes in the shared proportion of transboundary stocks include strengthening of current cooperative mechanisms and the consideration of side payments (including non-monetary arrangements) (Miller et al., 2013; Tunca, 2019), increased international cooperation (Miller et al., 2013) and management rules that capture distributional shifts (Palacios-Abrantes, Sumaila, et al., 2020; Pinsky et al., 2018). Some examples of current treaties that have adapted some of these strategies include: the Pacific Salmon Treaty between Canada and the U.S. to manage Pacific salmon (Oncorhynchus sp.) and which has a conservation fund functioning as a side payment (Miller et al., 2013), the International Pacific Halibut Commission between Canada and the United States, which as the name implies manages Pacific halibut (*Hippoglossus stenolepis*) and allocates quota based on the stock’s yearly distribution (Palacios-Abrantes, Sumaila, et al., 2020); and the Parties to the Nauru Agreement (PNA) an Oceania subregional agreement among 8 Pacific Island countries and Tokelau, which collectively manage the largest tuna fishery in the world, controlling around 50% of the global supply of skipjack tuna (*Katsuwonus pelamis*). The PNA manage fishing effort under a so-called Vessel Day Scheme, an adaptive and equitable system which accounts for shifts in stocks and catches as a result of climate variability (Bahri et al., 2021). Quota allocation methods based on a stock’s current distribution and/or according to fixed-historical proportions like that of the European Union need to evolve to be more agile (Baudron et al., 2020) and potentially move towards a dynamic method or a combination of both (Palacios-Abrantes, Sumaila, et al., 2020; Sumaila et al., 2020).

Even management plans that take into consideration such strategies might not be fully prepared for the consequences of shifting transboundary stocks (Engler, 2020; Koubrak & VanderZwaag, 2020; Pinsky et al., 2018). For example, species’ shifts within the PNA area will likely also have to deal with stocks expanding to new jurisdictions, issue that the NEAFC is currently facing regarding Atlantic mackerel (Pinsky et al., 2018; Spijkers & Boonstra, 2017). Moreover, the transition from historic to dynamic allocations can find strong resistance from stakeholders “losing” benefits from a fishery they have historically been entitled to. Finally, changes in policy can be sluggish (e.g., it took the PST about 10 years to agree on a conservation fund) compared to the timeframe over which species shift (Pinsky & Fogarty, 2012). This is specifically true as in many cases we lack standardized data to track shifts of species across international jurisdictions (Maureaud et al., 2021) as these might already be happening. Addressing shifting transboundary stocks is also challenged by an overall lack of knowledge of how many international agreements exist or what transboundary stocks are jointly managed. Recent efforts looking at 127 international fisheries plans found that they all lacked direct actions to address topics of climate change or species range shifts across jurisdictions (Oremus et al., 2020). Having a better understanding of what stocks are managed as transboundary and under what rules would be an important complement to our study and an important step towards the sustainability of international stocks. (Oremus et al., 2020).

### Potential drivers of shift and managing an uncertain future

The early projected range shifts of large number of transboundary stocks can be partially attributed to the parallel global emergence of several ocean variables from historical variability. For example, sea surface temperature (SST) is projected to increase beyond historical variability by 2030 in 50 to 70% of the global ocean (Frölicher et al., 2016; Mahlstein et al., 2011; Rodgers et al., 2015). Results from multiple marine ecosystem models show that SST and primary production (NPP) are the main indicators of species distributional changes across ocean basins (Bryndum-Buchholz et al., 2019; Lotze et al., 2019). Specifically, in the species distribution model used here (see methods), SST is the main environmental driver of biomass changes in polar, tropical and upwelling ecosystems, while NPP drives temperate regions’ results. The combination of the SST emergence pattern and model characteristics could be partially responsible for the early distributional shift of transboundary stocks in the tropics, where marine species live close to their thermal tolerance, making them highly vulnerable to warming waters (IPCC, 2019), while also explaining the later, and sometimes non-existent, shifts at higher latitudes (Fig. 3B). Different levels of uncertainty exist in the projected emergence of environmental variables mainly related to *i*) climate change scenarios and *ii*) model structure (Frölicher et al., 2016). First, our analysis is based on a high emission scenario (RCP 8.5) representing an “extreme” case of shifts in the shared distribution of transboundary species. Thus, any mitigation efforts could result in a substantial delay in distribution shifts as environmental signals are responsive to mitigation, at least after the middle of the 21^*st*^ century (Frölicher et al., 2016). Second, a substantial source of uncertainty is related to model selection for both climate change and fish distribution. This is specifically important for early time periods (e.g., 2016-2035) where the uncertainty related to model selection, including the parameterization of poorly understood processes that regulate NPP changes, is often larger than the uncertainty stemming from the climate change scenario (Frölicher et al., 2016). While such uncertainty could potentially be reduced by incorporating ensemble simulations from a range of different ESMs, these are computationally expensive simulations (Frölicher et al., 2009; Frölicher et al., 2016; Rodgers et al., 2015) that are only just becoming available (Deser et al., 2020) and cannot address limitations of process-level understanding. It is unlikely that these complex uncertainties will adequately be addressed in the coming decade. This is critical for the tropics, where the shared distribution of transboundary stocks is expected to shift the earliest, and the response of the base of the food web to climate change is most uncertain (Kwiatkowski et al., 2020; Tagliabue et al., 2020). These uncertainties also affect the projection of species distributions (Bryndum-Buchholz et al., 2019; Lotze et al., 2019). Overall, multiple upper trophic level models present overall agreement in terms of directional change of fish biomass, but variability in the magnitude of that change (Lotze et al., 2019). Further research that reduces uncertainty in the NPP response at the base of ocean food webs, alongside large ensemble simulations of multiple species distribution models could lead to smaller structural uncertainty of fish and fisheries models (Bryndum-Buchholz et al., 2019). Another important source of uncertainty in this study is the utilization of political boundaries (EEZ) to define stocks. While this method might define some stocks that do not necessarily align with biologically-defined subpopulations within an EEZ, in many EEZs, fisheries are often managed at the species level (MAP, 2017) and sub-populations are potentially interconnected (Ramesh et al., 2019), thus, providing additional ecological ground for our analysis (Dunn et al., 2019). Reproducing our analysis regionally, where spatially explicit stock data is available would allow to generate better constrained results and potentially identify different types of shared stock shifts at metapopulation level (Archambault et al., 2016; Link et al., 2010). Addressing these uncertainties systematically can serve as a roadmap for future studies to provide additional information to inform policy towards sustainable and equitable international fisheries management under climate change.

The global community has set the ambitious goal of managing all fisheries sustainably (SDG 14 - Life below water) by 2030; achieving this goal would have clear benefits for several other societal goals (Singh et al., 2017; United Nations, 2018). Developing anticipatory policies to deal with shifting transboundary stocks is key to achieving the SDGs and ensuring effective governance of the world’s fisheries (Oremus et al., 2020; Pecl et al., 2017; Pinsky et al., 2018). Here, we developed an approach to inform the sustainable management and governance of transboundary fisheries in a changing world (Link et al., 2010). First, we identified the transboundary stocks that would likely see shifts in their shared distribution compared to their historical average and the year in which such shifts would occur. Second, we estimated the intensity of such change. While future studies, specifically at more localized scale, will provide valuable nuance in designing effective policies, our results provide an important baseline on which to build when preparing ocean governance for shifting transboundary stocks (Palacios-Abrantes, Reygondeau, et al., 2020; Pinsky et al., 2018). Our findings emphasize recent calls for the urgent adoption of measures in support of more adaptive, flexible fisheries management and governance to support resilient fisheries and durable management systems (Oremus et al., 2020; Pinsky et al., 2018; Pinsky et al., 2020).

