## Supplemental material for "Timing and magnitude of climate driven range shifts in transboundary fish stocks challenge their management"

#### Figures

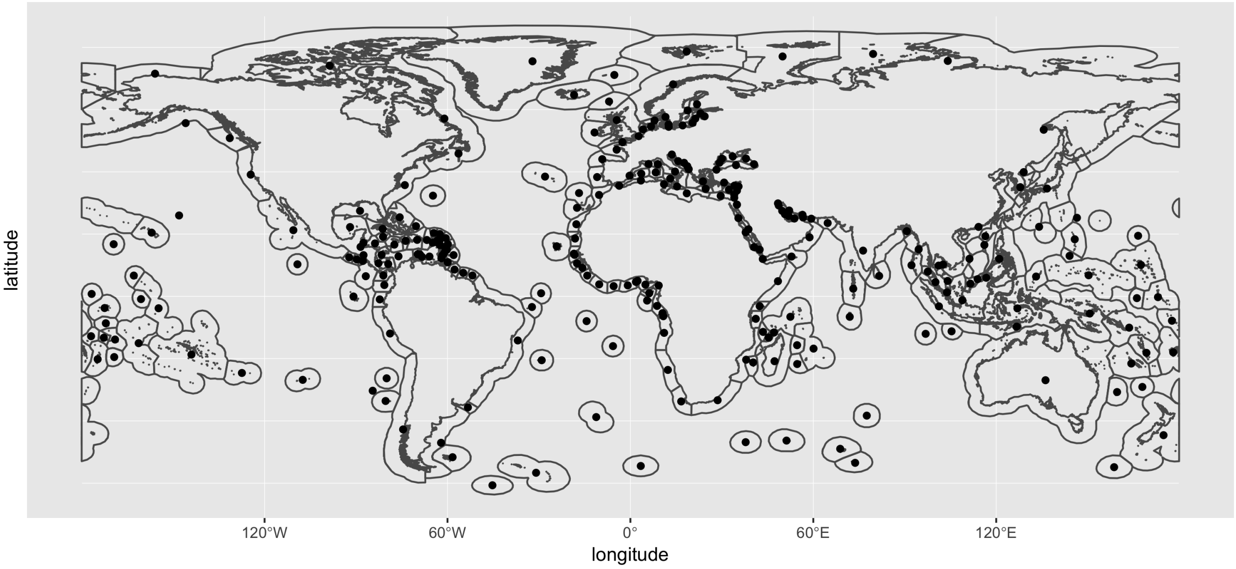


Figure S 1: World Exclusive Economic Zones used in this study as divided by the Sea Around Us and their estimated centroids (black points).

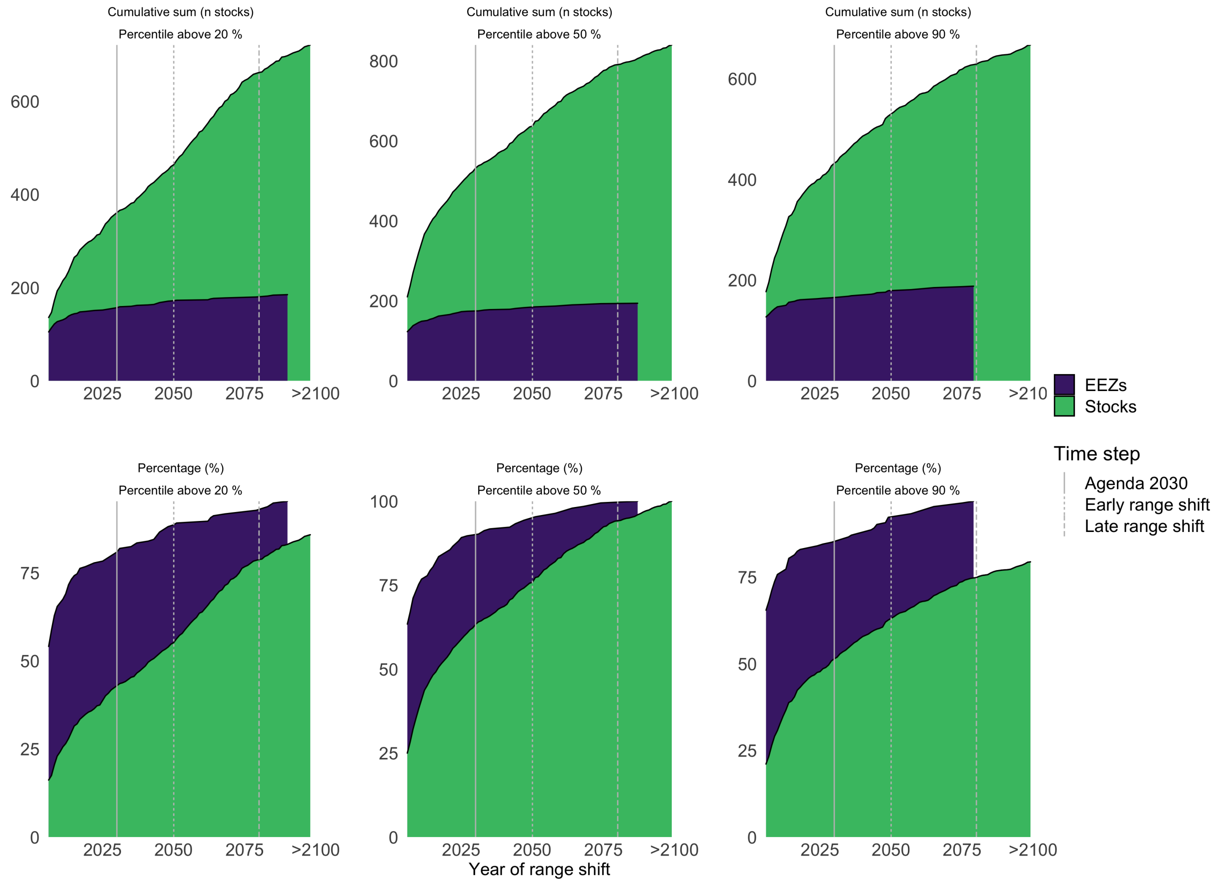


Figure S 2: Average year of range shift by Exclusive Economic Zone (EEZ) and transboundary stock using different levels of abundance percentile to determine the stock's centroid. Top Row: results in absolute numbers. Bottom row: results expressed as a percentage of the total emerging stocks and EEZs with emerging stock.

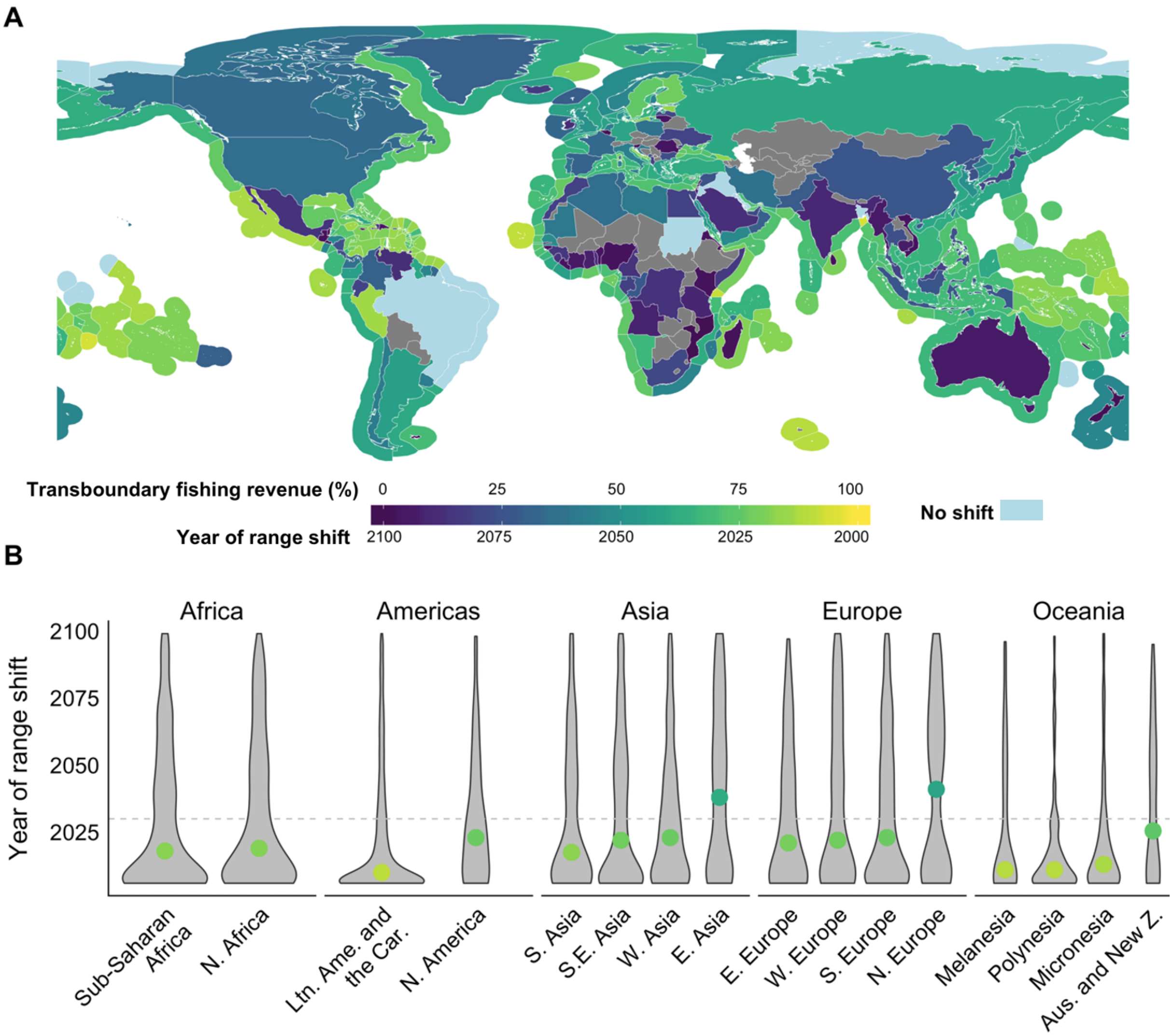


Figure S 3: Year of change in the shared distribution of 5,745 transboundary stocks using a one standard deviation threshold. A) Land polygons show the contribution of shifting stocks to a country or territory's total fishing revenue from transboundary stocks. Exclusive Economic Zone polygons display average year of range shifts within them. EEZs with no distributional shift between 2006 and 2100 are represented in aqua color. B) Year of shared distribution shifts by UN sub-regions. Points color coded by the median year of range shift. Horizontal dashed line represents the year 2030 – the Sustainable Development Agenda deadline. N = North, S = South, W = West and E = East. Ltn. Ame. and the Car. = Latin America and the Caribbean. Aus and New Z. = Australia and New Zealand.

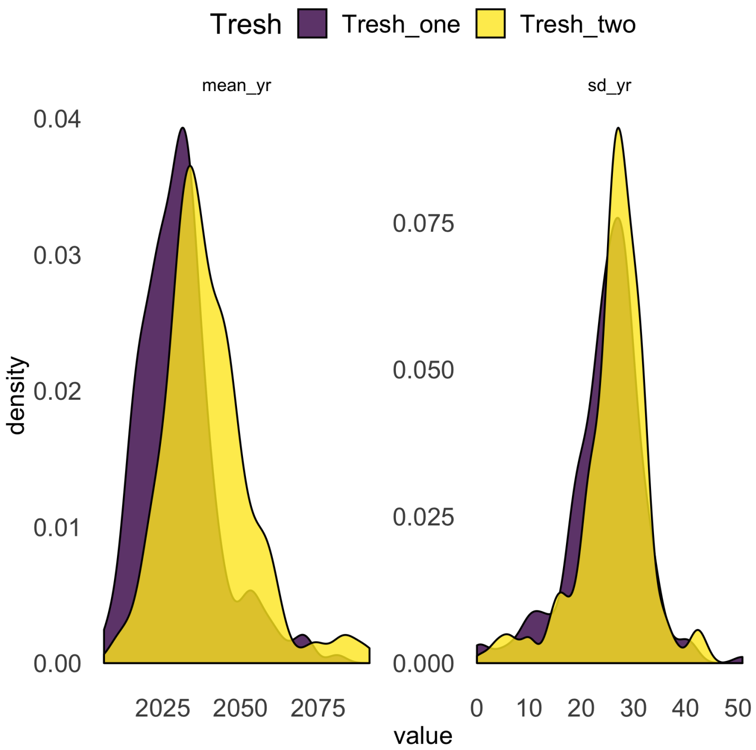


Figure S 4: Average distribution of transboundary stock's year of change in their shared distribution. Left panel is the average year across stocks and right panel the standard deviation. Yellow represents a two standard deviation distance from the historical average and purple represents a one standard deviation distance from the historical average.

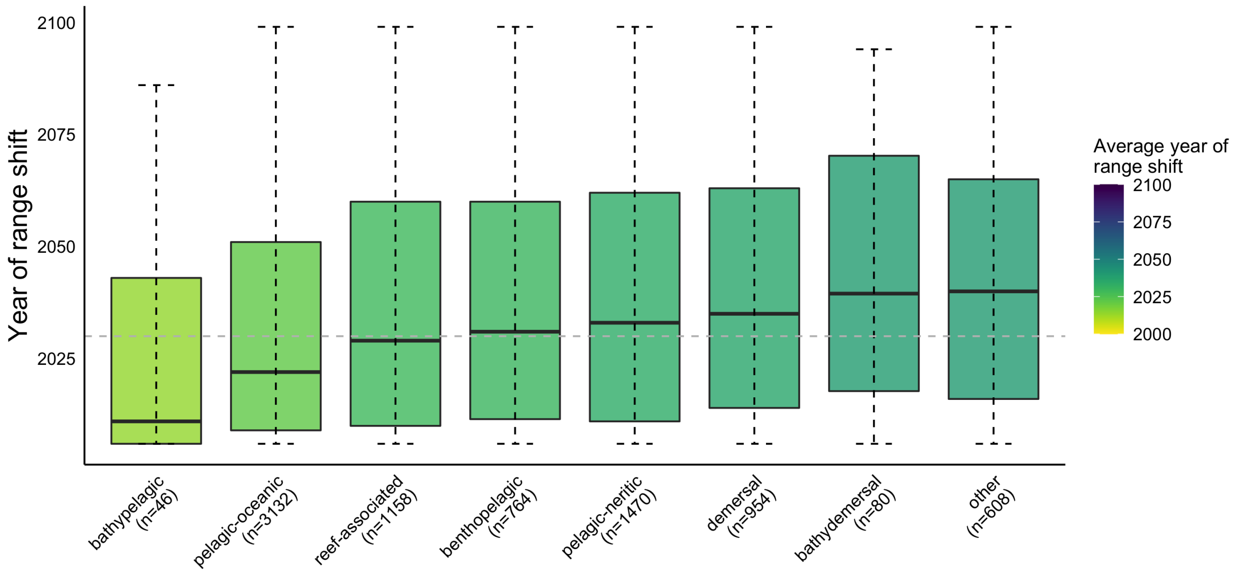


Figure S 5: Comparison of shifts in the shared distribution of transboundary stocks by species' habitat association. Classification is based on the habitat preference information obtained from FishBase and SeaLifeBase (see Table S1). The category "Other" consists of species that have no "habitat preference" classification. The number of stocks included in this analysis for each habitat type is noted in parentheses. Dashed whiskers represent the 1.5* interquartile range. Box represents interquartile range as distance between first and third quartiles. Solid line represents the median time of emergence per habitat preference.

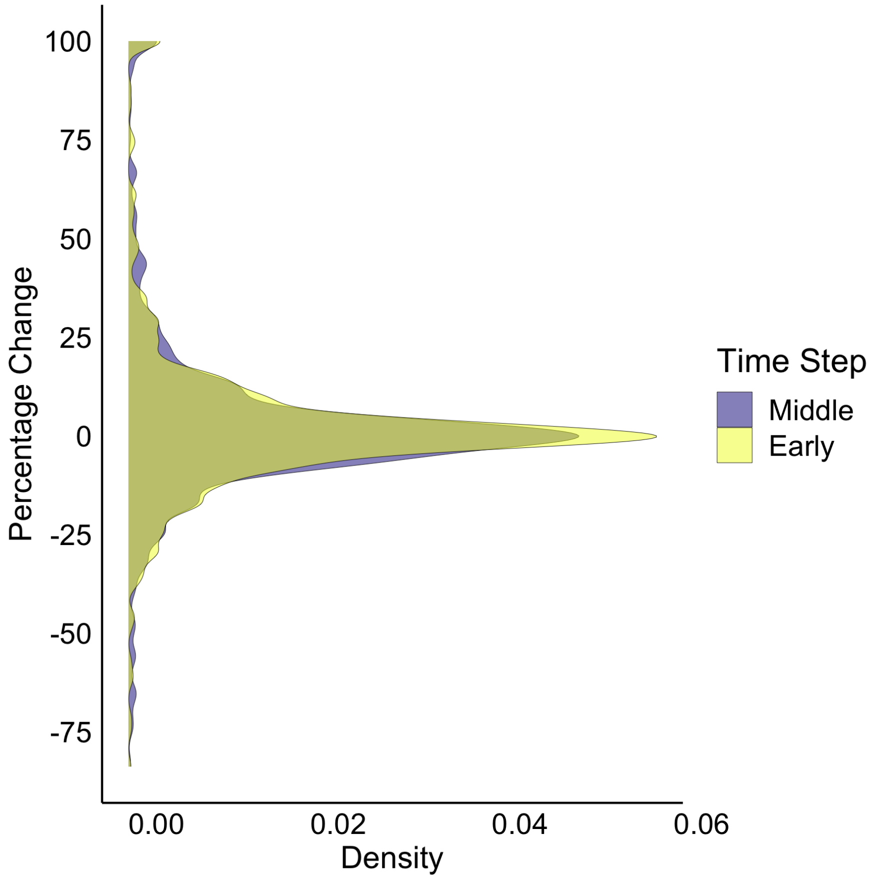


Figure S 6: Distribution of changes in transboundary stock proportion per stock and neighboring Exclusive Economic Zones. Yellow color represents changes by early, 2030 (2021-2040) and purple by mid 2050 (2041-60) 21st century relative to 1951-2005.

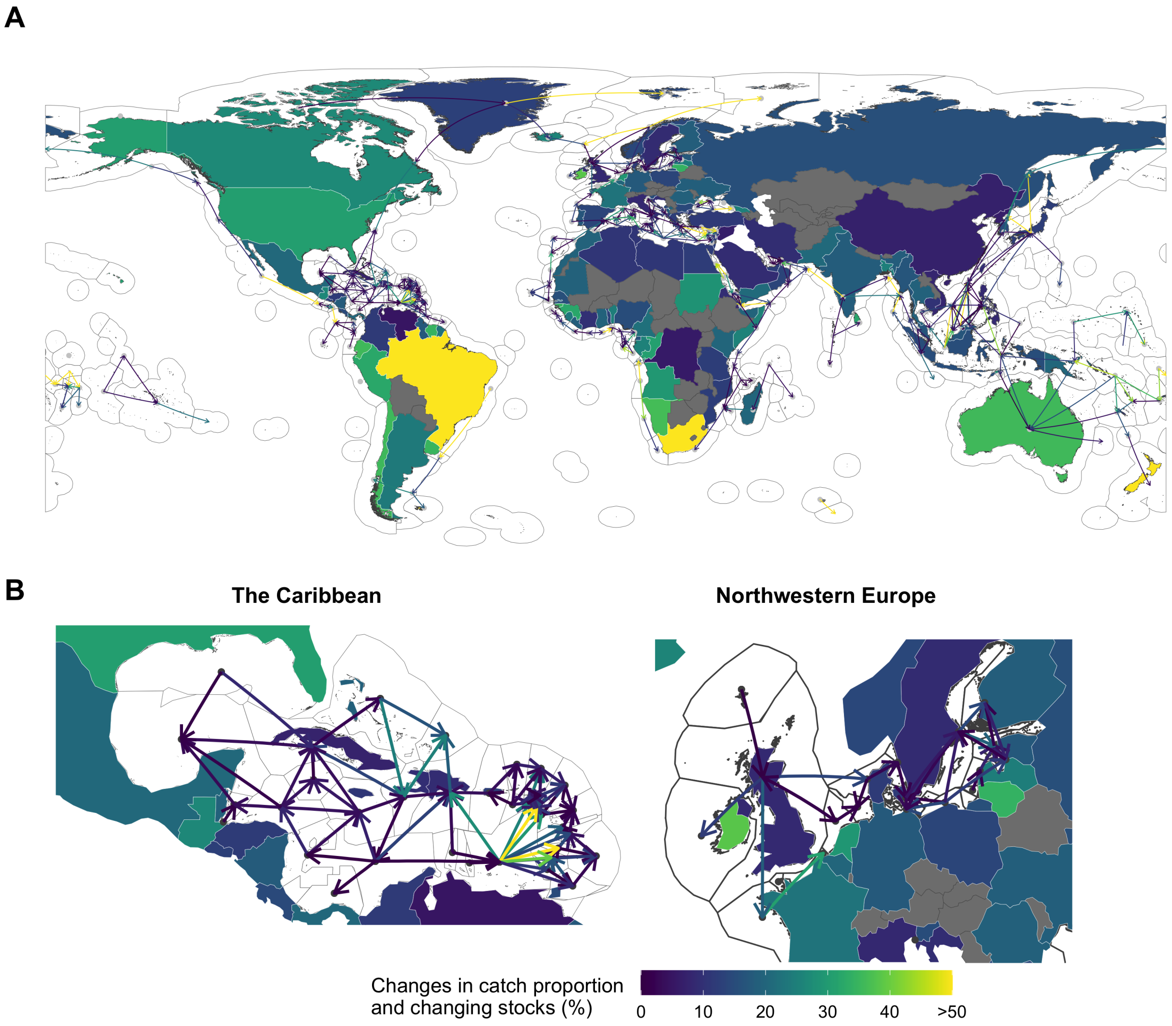


Figure S 7: Changes in stock proportion of neighboring Exclusive Economic Zones by 2050 (2041-2060) relative to 1951-2005. Lines represent the average change in transboundary stock share ratio with arrows going from EEZ with decreasing stock share (point) to those gaining shares (arrowhead). Land polygons represent the percentage of stocks that are projected to change their stock share ratio beyond the identified threat point with higher gains identified in warmer colors. Panel B highlights changes for the Caribbean and Northwestern Europe.

### Tables

Table S 1: Definition of each habitat preference according to FishBase

| Habitat preference | Description |
|---|---|
| pelagic-neritic | The shallow pelagic zone over the continental shelf; nearshore ocean ecosystems; i.e., those associated with the coasts because the waters are overlying continental shelves and/or the waters are < 200 m deep in areas of coastal submarine slopes |
| reef-associated | A largely consolidated wave resistant feature; the upper surface is within 0-20 m of the ocean surface; in the tropics, the upper portion is typically composed mainly of living and non-living remains of coral and coralline algae; rock lying at or near the surface and which may pose a danger to navigation |
| bathydemersal | Living and feeding on the bottom below 200 m |
| benthopelagic | Living and feeding near the bottom as well as in midwaters or near the surface. Feeding on benthic as well as free swimming organisms. Many freshwater fish are opportunistic feeders that forage on the bottom as well as in midwater and near the surface, also pertaining to forms which hover or swim just over the floor of the sea, e.g. Halosauridae, Macrouridae, Moridae, Brotulidae; the depth zone about 100 metres off the bottom at all depths below the edge of the continental shelf |
| demersal | Sinking to or lying on the bottom; living on or near the bottom and feeding on benthic organisms |
| bathypelagic | Region of the oceanic zone between 1,000 m to 4,000 m; between the mesopelagic layer above and the abyssopelagic layer below. Living or feeding in open waters at depths between 1,000 and 4,000 m. In FishBase this term is used to include the depth range from 200 m to the bottom and thus the zones mesopelagic, bathypelagic and abyssopelagic |
| pelagic-oceanic | Living and feeding in the open sea; associated with the surface or middle depths of a body of water; free swimming in the seas, oceans or open waters; not in association with the bottom. Many pelagic fish feed on plankton. In FishBase, referring to surface or mid water from 0 to 200 m depth. |

Table S 2: Statistical results for multiple comparison test after Kruskal-Wallis on time of emergence across United Nations sub regions

| Sub region comparrison | Significan level | Observation difference | Critical difference | K-W Difference |
|---|---|---|---|---|
| bathydemersal-bathypelagic | 0.05 | 1618.20177 | 1347.2420 | TRUE |
| bathydemersal-benthopelagic | 0.05 | 769.28313 | 841.6236 | FALSE |
| bathydemersal-demersal | 0.05 | 664.41978 | 832.8192 | FALSE |
| bathydemersal-pelagic | 0.05 | 784.99848 | 1851.4997 | FALSE |
| bathydemersal-pelagic-neritic | 0.05 | 1040.43584 | 823.9146 | TRUE |
| bathydemersal-pelagic-oceanic | 0.05 | 1633.56909 | 810.1675 | TRUE |
| bathydemersal-reef-associated | 0.05 | 1353.67920 | 829.8040 | TRUE |
| bathypelagic-benthopelagic | 0.05 | 848.91864 | 1119.7065 | FALSE |
| bathypelagic-demersal | 0.05 | 953.78199 | 1113.1039 | FALSE |
| bathypelagic-pelagic | 0.05 | 833.20329 | 1993.3548 | FALSE |
| bathypelagic-pelagic-neritic | 0.05 | 577.76593 | 1106.4572 | FALSE |
| bathypelagic-pelagic-oceanic | 0.05 | 15.36731 | 1096.2590 | FALSE |
| bathypelagic-reef-associated | 0.05 | 264.52257 | 1110.8497 | FALSE |
| benthopelagic-demersal | 0.05 | 104.86335 | 363.6887 | FALSE |
| benthopelagic-pelagic | 0.05 | 15.71535 | 1693.1429 | FALSE |
| benthopelagic-pelagic-neritic | 0.05 | 271.15271 | 342.8074 | FALSE |
| benthopelagic-pelagic-oceanic | 0.05 | 864.28595 | 308.3068 | TRUE |
| benthopelagic-reef-associated | 0.05 | 584.39607 | 356.7300 | TRUE |
| demersal-pelagic | 0.05 | 120.57870 | 1688.7837 | FALSE |
| demersal-pelagic-neritic | 0.05 | 376.01606 | 320.5846 | TRUE |
| demersal-pelagic-oceanic | 0.05 | 969.14931 | 283.3914 | TRUE |
| demersal-reef-associated | 0.05 | 689.25942 | 335.4309 | TRUE |
| pelagic-pelagic-neritic | 0.05 | 255.43736 | 1684.4102 | FALSE |
| pelagic-pelagic-oceanic | 0.05 | 848.57060 | 1677.7288 | FALSE |
| pelagic-reef-associated | 0.05 | 568.68071 | 1687.2988 | FALSE |
| pelagic-neritic-pelagic-oceanic | 0.05 | 593.13324 | 256.0431 | TRUE |
| pelagic-neritic-reef-associated | 0.05 | 313.24335 | 312.6680 | TRUE |
| pelagic-oceanic-reef-associated | 0.05 | 279.88989 | 274.4039 | TRUE |

Table S 3: Statistical results for multiple comparison test after Kruskal-Wallis on time of emergence across United Nations sub regions

| habitat association comparrison | Significan level | Observation difference | Critical difference | K-W Difference |
|---|---|---|---|---|
| Australia and New Zealand-Eastern Asia | 0.05 | 715.011934 | 1465.9298 | FALSE |
| Australia and New Zealand-Eastern Europe | 0.05 | 564.958758 | 1605.2391 | FALSE |
| Australia and New Zealand-Latin America and the Caribbean | 0.05 | 1569.939379 | 1381.8702 | TRUE |
| Australia and New Zealand-Melanesia | 0.05 | 1127.076002 | 1558.6091 | FALSE |
| Australia and New Zealand-Micronesia | 0.05 | 1284.742986 | 1954.7306 | FALSE |
| Australia and New Zealand-Northern Africa | 0.05 | 367.027459 | 1426.8945 | FALSE |
| Australia and New Zealand-Northern America | 0.05 | 319.846054 | 1634.3439 | FALSE |
| Australia and New Zealand-Northern Europe | 0.05 | 597.982863 | 1487.1176 | FALSE |
| Australia and New Zealand-Polynesia | 0.05 | 1711.663455 | 1611.5762 | TRUE |
| Australia and New Zealand-South-eastern Asia | 0.05 | 248.225363 | 1397.9867 | FALSE |
| Australia and New Zealand-Southern Asia | 0.05 | 522.496243 | 1526.7065 | FALSE |
| Australia and New Zealand-Southern Europe | 0.05 | 20.481074 | 1397.5455 | FALSE |
| Australia and New Zealand-Sub-Saharan Africa | 0.05 | 561.927280 | 1398.5285 | FALSE |
| Australia and New Zealand-Western Asia | 0.05 | 202.306373 | 1440.4445 | FALSE |
| Australia and New Zealand-Western Europe | 0.05 | 10.591741 | 1495.5026 | FALSE |
| Eastern Asia-Eastern Europe | 0.05 | 1279.970692 | 995.4690 | TRUE |
| Eastern Asia-Latin America and the Caribbean | 0.05 | 2284.951313 | 568.9736 | TRUE |
| Eastern Asia-Melanesia | 0.05 | 1842.087936 | 918.3835 | TRUE |
| Eastern Asia-Micronesia | 0.05 | 1999.754921 | 1495.0377 | TRUE |
| Eastern Asia-Northern Africa | 0.05 | 1082.039393 | 670.9648 | TRUE |
| Eastern Asia-Northern America | 0.05 | 1034.857988 | 1041.7513 | FALSE |
| Eastern Asia-Northern Europe | 0.05 | 117.029071 | 791.0022 | FALSE |
| Eastern Asia-Polynesia | 0.05 | 2426.675389 | 1005.6558 | TRUE |
| Eastern Asia-South-eastern Asia | 0.05 | 963.237297 | 607.0687 | TRUE |
| Eastern Asia-Southern Asia | 0.05 | 1237.508177 | 863.1329 | TRUE |
| Eastern Asia-Southern Europe | 0.05 | 735.493008 | 606.0521 | TRUE |
| Eastern Asia-Sub-Saharan Africa | 0.05 | 1276.939214 | 608.3155 | TRUE |
| Eastern Asia-Western Asia | 0.05 | 917.318307 | 699.3183 | TRUE |
| Eastern Asia-Western Europe | 0.05 | 704.420193 | 806.6559 | FALSE |
| Eastern Europe-Latin America and the Caribbean | 0.05 | 1004.980621 | 866.9333 | TRUE |
| Eastern Europe-Melanesia | 0.05 | 562.117244 | 1127.5064 | FALSE |
| Eastern Europe-Micronesia | 0.05 | 719.784229 | 1631.8640 | FALSE |
| Eastern Europe-Northern Africa | 0.05 | 197.931299 | 937.0358 | FALSE |

Table S 3: Statistical results for multiple comparison test after Kruskal-Wallis on time of emergence across United Nations sub regions *(continued)*

| habitat association comparrison | Significan level | Observation difference | Critical difference | K-W Difference |
|---|---|---|---|---|
| Eastern Europe-Northern America | 0.05 | 245.112704 | 1230.0765 | FALSE |
| Eastern Europe-Northern Europe | 0.05 | 1162.941621 | 1026.4146 | TRUE |
| Eastern Europe-Polynesia | 0.05 | 1146.704697 | 1199.6607 | FALSE |
| Eastern Europe-South-eastern Asia | 0.05 | 316.733395 | 892.3984 | FALSE |
| Eastern Europe-Southern Asia | 0.05 | 42.462515 | 1082.9778 | FALSE |
| Eastern Europe-Southern Europe | 0.05 | 544.477684 | 891.7071 | FALSE |
| Eastern Europe-Sub-Saharan Africa | 0.05 | 3.031478 | 893.2469 | FALSE |
| Eastern Europe-Western Asia | 0.05 | 362.652385 | 957.5429 | FALSE |
| Eastern Europe-Western Europe | 0.05 | 575.550499 | 1038.5259 | FALSE |
| Latin America and the Caribbean-Melanesia | 0.05 | 442.863377 | 777.2021 | FALSE |
| Latin America and the Caribbean-Micronesia | 0.05 | 285.196393 | 1412.7110 | FALSE |
| Latin America and the Caribbean-Northern Africa | 0.05 | 1202.911920 | 459.1390 | TRUE |
| Latin America and the Caribbean-Northern America | 0.05 | 1250.093325 | 919.7069 | TRUE |
| Latin America and the Caribbean-Northern Europe | 0.05 | 2167.922242 | 621.5299 | TRUE |
| Latin America and the Caribbean-Polynesia | 0.05 | 141.724076 | 878.6116 | FALSE |
| Latin America and the Caribbean-South-eastern Asia | 0.05 | 1321.714017 | 359.3708 | TRUE |
| Latin America and the Caribbean-Southern Asia | 0.05 | 1047.443136 | 711.0649 | TRUE |
| Latin America and the Caribbean-Southern Europe | 0.05 | 1549.458305 | 357.6508 | TRUE |
| Latin America and the Caribbean-Sub-Saharan Africa | 0.05 | 1008.012099 | 361.4728 | TRUE |
| Latin America and the Caribbean-Western Asia | 0.05 | 1367.633007 | 499.6608 | TRUE |
| Latin America and the Caribbean-Western Europe | 0.05 | 1580.531120 | 641.3335 | TRUE |
| Melanesia-Micronesia | 0.05 | 157.666984 | 1586.0170 | FALSE |
| Melanesia-Northern Africa | 0.05 | 760.048543 | 854.6964 | FALSE |
| Melanesia-Northern America | 0.05 | 807.229948 | 1168.5709 | FALSE |
| Melanesia-Northern Europe | 0.05 | 1725.058865 | 951.8386 | TRUE |
| Melanesia-Polynesia | 0.05 | 584.587453 | 1136.5103 | FALSE |
| Melanesia-South-eastern Asia | 0.05 | 878.850640 | 805.5089 | TRUE |
| Melanesia-Southern Asia | 0.05 | 604.579759 | 1012.5762 | FALSE |
| Melanesia-Southern Europe | 0.05 | 1106.594928 | 804.7431 | TRUE |
| Melanesia-Sub-Saharan Africa | 0.05 | 565.148722 | 806.4489 | FALSE |
| Melanesia-Western Asia | 0.05 | 924.769629 | 877.1307 | TRUE |
| Melanesia-Western Europe | 0.05 | 1137.667743 | 964.8865 | TRUE |

Table S 3: Statistical results for multiple comparison test after Kruskal-Wallis on time of emergence across United Nations sub regions *(continued)*

| habitat association comparrison | Significan level | Observation difference | Critical difference | K-W Difference |
|---|---|---|---|---|
| Micronesia-Northern Africa | 0.05 | 917.715528 | 1456.7825 | FALSE |
| Micronesia-Northern America | 0.05 | 964.896933 | 1660.5021 | FALSE |
| Micronesia-Northern Europe | 0.05 | 1882.725850 | 1515.8186 | TRUE |
| Micronesia-Polynesia | 0.05 | 426.920469 | 1638.0980 | FALSE |
| Micronesia-South-eastern Asia | 0.05 | 1036.517624 | 1428.4796 | FALSE |
| Micronesia-Southern Asia | 0.05 | 762.246743 | 1554.6768 | FALSE |
| Micronesia-Southern Europe | 0.05 | 1264.261912 | 1428.0479 | FALSE |
| Micronesia-Sub-Saharan Africa | 0.05 | 722.815706 | 1429.0099 | FALSE |
| Micronesia-Western Asia | 0.05 | 1082.436614 | 1470.0570 | FALSE |
| Micronesia-Western Europe | 0.05 | 1295.334727 | 1524.0457 | FALSE |
| Northern Africa-Northern America | 0.05 | 47.181405 | 986.0646 | FALSE |
| Northern Africa-Northern Europe | 0.05 | 965.010322 | 716.0741 | TRUE |
| Northern Africa-Polynesia | 0.05 | 1344.635996 | 947.8508 | TRUE |
| Northern Africa-South-eastern Asia | 0.05 | 118.802096 | 505.5790 | FALSE |
| Northern Africa-Southern Asia | 0.05 | 155.468784 | 795.0321 | FALSE |
| Northern Africa-Southern Europe | 0.05 | 346.546385 | 504.3579 | FALSE |
| Northern Africa-Sub-Saharan Africa | 0.05 | 194.899821 | 507.0753 | FALSE |
| Northern Africa-Western Asia | 0.05 | 164.721086 | 613.2893 | FALSE |
| Northern Africa-Western Europe | 0.05 | 377.619200 | 733.3290 | FALSE |
| Northern America-Northern Europe | 0.05 | 917.828917 | 1071.3609 | FALSE |
| Northern America-Polynesia | 0.05 | 1391.817401 | 1238.3349 | TRUE |
| Northern America-South-eastern Asia | 0.05 | 71.620691 | 943.7490 | FALSE |
| Northern America-Southern Asia | 0.05 | 202.650189 | 1125.6679 | FALSE |
| Northern America-Southern Europe | 0.05 | 299.364980 | 943.0954 | FALSE |
| Northern America-Sub-Saharan Africa | 0.05 | 242.081227 | 944.5515 | FALSE |
| Northern America-Western Asia | 0.05 | 117.539681 | 1005.5724 | FALSE |
| Northern America-Western Europe | 0.05 | 330.437794 | 1082.9697 | FALSE |
| Northern Europe-Polynesia | 0.05 | 2309.646318 | 1036.2972 | TRUE |
| Northern Europe-South-eastern Asia | 0.05 | 846.208226 | 656.5828 | TRUE |
| Northern Europe-Southern Asia | 0.05 | 1120.479106 | 898.6473 | TRUE |
| Northern Europe-Southern Europe | 0.05 | 618.463937 | 655.6429 | FALSE |
| Northern Europe-Sub-Saharan Africa | 0.05 | 1159.910143 | 657.7356 | TRUE |

Table S 3: Statistical results for multiple comparison test after Kruskal-Wallis on time of emergence across United Nations sub regions *(continued)*

| habitat association comparrison | Significan level | Observation difference | Critical difference | K-W Difference |
|---|---|---|---|---|
| Northern Europe-Western Asia | 0.05 | 800.289236 | 742.7076 | TRUE |
| Northern Europe-Western Europe | 0.05 | 587.391122 | 844.5485 | FALSE |
| Polynesia-South-eastern Asia | 0.05 | 1463.438092 | 903.7477 | TRUE |
| Polynesia-Southern Asia | 0.05 | 1189.167212 | 1092.3488 | TRUE |
| Polynesia-Southern Europe | 0.05 | 1691.182381 | 903.0651 | TRUE |
| Polynesia-Sub-Saharan Africa | 0.05 | 1149.736175 | 904.5856 | TRUE |
| Polynesia-Western Asia | 0.05 | 1509.357082 | 968.1289 | TRUE |
| Polynesia-Western Europe | 0.05 | 1722.255196 | 1048.2944 | TRUE |
| South-eastern Asia-Southern Asia | 0.05 | 274.270880 | 741.8995 | FALSE |
| South-eastern Asia-Southern Europe | 0.05 | 227.744289 | 415.5907 | FALSE |
| South-eastern Asia-Sub-Saharan Africa | 0.05 | 313.701918 | 418.8844 | FALSE |
| South-eastern Asia-Western Asia | 0.05 | 45.918990 | 542.6440 | FALSE |
| South-eastern Asia-Western Europe | 0.05 | 258.817103 | 675.3592 | FALSE |
| Southern Asia-Southern Europe | 0.05 | 502.015169 | 741.0679 | FALSE |
| Southern Asia-Sub-Saharan Africa | 0.05 | 39.431037 | 742.9200 | FALSE |
| Southern Asia-Western Asia | 0.05 | 320.189870 | 819.1022 | FALSE |
| Southern Asia-Western Europe | 0.05 | 533.087984 | 912.4561 | FALSE |
| Southern Europe-Sub-Saharan Africa | 0.05 | 541.446206 | 417.4097 | TRUE |
| Southern Europe-Western Asia | 0.05 | 181.825299 | 541.5064 | FALSE |
| Southern Europe-Western Europe | 0.05 | 31.072815 | 674.4456 | FALSE |
| Sub-Saharan Africa-Western Asia | 0.05 | 359.620908 | 544.0383 | FALSE |
| Sub-Saharan Africa-Western Europe | 0.05 | 572.519021 | 676.4801 | FALSE |
| Western Asia-Western Europe | 0.05 | 212.898113 | 759.3574 | FALSE |
